# Damping nonlinearity in agarose hydrogels under relative humidity: balancing network stiffness and energy dissipation

**DOI:** 10.64898/2026.05.02.722420

**Authors:** Ikenna Ojoboh, Manuel Dedola, Katherine Nelms, Charles de Kergariou, Ibrahim Patrick, Ludovico Cademartiri, James P. K. Armstrong, Adam W. Perriman, Fabrizio Scarpa

## Abstract

Sustainable, biodegradable elastomers are needed to replace fossil-based alternatives and reduce the environmental impact of traditional vibration damping materials. We investigate agarose-based hydrogels as eco-friendly vibration absorbers, examining the combined effects of polymer concentration (1–7 wt%), relative humidity (55–98%), and mechanical pre-stress on their dynamic mechanical properties. Frequency-dependent viscoelastic and vibration transmissibility tests, supported by Gaussian Process Regression (GPR), reveal that increasing agarose concentration enhances the storage modulus (*E*^*′*^) by over an order of magnitude, reaching *∼*5 MPa depending on humidity and applied prestress. Remarkably, the damping efficiency—characterised by the loss factor (*tan*(*δ*))—exhibits a highly non-monotonic trend. Maximum energy dissipation is observed at intermediate network densities, with tan(*δ*) up to 0.21 and a loss modulus of *∼* 515 kPa at 5 w% and 75% relative humidity, comparable to synthetic elastomers. GPR analysis shows that prestress controls nonlinear stiffening and transmissibility resonance behavior, while shifting peak damping from 5 wt% to 1 wt% agarose as prestress increases. These findings underscore the mechanical tunability and sustainability of agarose hydrogels, providing potential design guidance for biodegradable vibration mitigation materials.

## 1 Introduction

The push for sustainable materials has accelerated the search for biodegradable alternatives to replace petroleum-based elastomers in vibration control [1–3]. Conventional synthetic rubbers dissipate energy effectively via molecular viscoelasticity, but their environmental persistence drives demand for renewable soft materials with matching performance [2]. In this context, hydrogels – three-dimensional polymer networks saturated with water-have recently emerged as promising candidates, combining intrinsic biocompatibility, high water content and highly tunable mechanical properties with significant viscous dissipation [3, 4]. The mechanical behavior of hydrogels stems from the balance between elastic energy storage and viscous dissipation driven by polymer chain mobility [5]. However, a fundamental trade-off remains: higher polymer concentration increases stiffness but hampers molecular mobility, reducing viscous dissipation—so stiffness and damping do not scale together across compositions [6–9].

Despite their success in biomedical and soft matter applications, hydrogels face key limitations—low stiffness, limited durability, and sensitivity to temperature, humidity, and mechanical pre-stress—that have restricted their adoption in mechanical engineering[10, 11]. Recent research has therefore focused on enhancing hydrogel performance through complex molecular and microstructure architectures, including double-network systems and composite formulations incorporating reinforcing fillers or fibres [12–14]. Alginate–poloxamer systems exploit the thermoresponsive micellisation of Poloxamer 407 to template network formation, yielding highly dissipative structures [2, 15]. Recent advances have demonstrated that doping these networks with natural reinforcements – such as sepiolite nanoclays and multiscale cactus fibers – can dramatically enhance dynamic stiffness and vibration damping, achieving loss factors exceeding 0.4. at frequencies below 300 Hz[16], a vibrational performance difficult to attain with classical fossil-based elastomers and polymer [17]. The significant vibration damping behavior observed in the latter hydrogel composite systems also stems from slip-stick friction mechanisms existing between the natural fiber reinforcements and the matrix [18, 19]. While these approaches achieve significant damping, their structural complexity often obscures the underlying structure–property relationships existing the hydrogel system. [20]

Agarose hydrogels, natural polysaccharides derived from red algae, are widely used in microbiology, food processing, and biotechnology. Their ability to form thermoreversible hydrogels — with tunable gelation kinetics and adaptive mechanical properties—drives their versatility[21, 22]. Agarose solutions undergo a rapid sol-gel transition upon cooling, forming a 3D network dictated by their molecular structure and physicochemical properties. Gelation occurs in two stages: initially, randomly distributed coils link via hydrogen bonds to form double-helical associations, followed by the aggregation (driven by hydrogen bond) of these helices into a rigid, three-dimensional network [23, 24]. The initial coil-to-helix transition that takes place during cooling can be described by a mean-field Zimm-Bragg approach [25–27].

Agarose’s controllable architecture makes it an ideal platform for isolating viscoelastic damping mechanisms. However, its nonlinear, time-dependent behavior—including mechanical memory and cyclic structural rearrangements—remains poorly understood, especially under realistic nonlinear loading conditions critical for advanced vibration mitigation.[28, 29] Moreover, key damping metrics-such as dynamic modulus (*E*^*′*^), loss modulus (*E* ^*”*^), and loss factor (*η* or tan *δ*) – are highly sensitive to subtle variations in concentration, hydration state and applied pre-stress, therefore necessitating systematic investigation. In this study, we attempt to address these challenges through a comprehensive experimental and modeling framework to characterize biodegradable agarose hydrogels (1-7 wt %) under conditions relevant to practical vibration load applications. Vibration transmissibility testing under broad-spectrum white noise excitation is employed here to isolate the damping behavior. This approach enables direct quantification of frequency-dependent storage modulus (*E*^*′*^), loss modulus (*E*^*′′*^), and loss factor (tan(*δ*)) across a wide parameter design space encompassing agarose concentration, relative humidity (55%-98%) and mechanical prestress. Our results show that while the storage modulus increases by over an order of magnitude with concentration—reaching up to *∼*5 MPa, depending on environmental conditions and applied load—the damping efficiency follows a highly non-monotonic trend. Experimental measurements identified peak energy dissipation at intermediate network densities, where tan(*δ*) reaches 0.21 (with a corresponding loss modulus of 515 kPa) at a 5 wt % concentration under 75% relative humidity, achieving values directly comparable to conventional synthetic elastomers [30]. To rationalize these complex interactions, Gaussian Process Regression (GPR) is employed to map the equivalent viscoelasticity measured using the vibration transmissibility tests across agarose concentrations, relative humidity values and prestress values exerted over the hydrogel samples. These predictive surfaces demonstrate that mechanical prestress primarily drives nonlinear stiffening and lowers resonance frequency, while also reshaping the damping landscape. As prestress increases, the optimal polymer concentration for maximum dissipation shifts—from 5 wt% at low prestress to just 1 wt% at high prestress. This predictive framework may enable the identification of material and environmental design parameters that maximize energy dissipation in these biodegradable soft materials.

## 2 Materials and Methods

### 2.1 Preparation of agarose hydrogels

#### 2.1.1 Synthesis

Agarose hydrogels with concentrations ranging from 1 to 7 wt% were prepared by dissolving related quantities of Agarose BP160-100 powder (Fisher Scientific, USA) in deionized water. The solutions were autoclaved at 120°C for 20 minutes to ensure complete dissolution, cooled to room temperature to allow initial gelation and subsequently refrigerated at 5°C.

#### 2.1.2 Casting and shaping

Prior to sample fabrication, the gels were reheated to *∼* 100°C until fully liquefied. The molten agarose was then cast into 3D-printed molds to form blocks of 30 *×* 30 *×* 15 mm^3^. Once solidified, the blocks were trimmed to provide a regular shape for the fitting in the vibration transmissibility rig (see Supporting information, Tab. S5,S6,S7). Samples were also prepared for quasi-static compression testing. For that case, the liquefied agarose was poured into Petri dishes, allowed to gel, and cored using a metal punch to yield cylindrical specimens (13 mm of diameter and 5 mm of height). Due to the soft nature of the hydrogels, slight height variations occurred. To ensure accurate mechanical data normalization, the exact height (*L*_0_) of each specimen was measured with a caliper immediately before testing (see Supporting Information, Tab. S2).

#### 2.1.3 Conditioning of the specimens

Hydrogel samples were conditioned at three relative humidity (RH) values using monitored environmental conditions and solution or salts [31]. The 55% RH was achieved in ambient controlled conditions, while the 75% and 98% with *NaCl* and *K*_2_*SO*_4_ solutions, respectively. Saturated salt solution were prepared with deionized water and excess salts, and stirred for 30 minutes to ensure saturation. The agarose hydrogels were placed within airtight plastic containers on platforms above the saturated salt solutions, sealed and refrigerated for 7 days to archive the targeted RH conditions.[32]

### 2.2 Mechanical and dynamics characterization

#### 2.2.1 Quasistatic compression testing

Quasistatic uniaxial compression tests were performed using an FMS-500 L2 Force Measurement System (Starrett, Australia) equipped with a 100 N load cell. The top platen was lowered until initial contact with the sample surface was detected. During the compression test, a quasistatic displacement rate of 0.6 mm*·*min^−1^ was applied. This rate corresponds to a strain of approximately *∼* 0.18 % *s*^−1^ which falls below the critical threshold of 0.7% *s*^−1^ identifyed by Ed-Dauby *et al* [33]. Operating within this range ensures a compressible poroelastic response, enabling interstitial fluid drainage during deformation.

#### 2.2.2 Vibration Transmissibility Testing

Vibration transmissibility tests were conducted using a dynamic shaker controlled via NI-DAQmx in MATLAB, interfaced with two data acquisition units (National Instruments USB-6211 and cDAQ-9178). The experimental apparatus consisted of a shaker powered by an LDS PA100E amplifier (gain = 5) and accelerometers (model 333 M07, PCB Piezotronics, USA) mounted on the rig’s base and top plates. Samples were centrally aligned, secured onto a rigid base plate, and prestressed by applying top masses (*M*) of 99 g, 146 g and 195 g. The masses were used to vary the resonant frequency of the transmissibility and therefore detect the variation of the dynamic/storage modulus and loss factor at those frequencies. The samples were subjected to white noise input at four acceleration amplitudes (0.1–0.625 m/s^2^) to assess responses linearity and characteristics. Input signals generated in MATLAB were amplified through the LDS PA100E amplifier and transmitted to the shaker. Base and sample vibrations were captured by PCB accelerometers acquired via NI USB-6211 and cDAQ-9178 data acquisition cards and units [2].

#### 2.2.3 Dynamic Mechanical Analyzer

Longitudinal DMA was performed on a TA Instruments DMA850 equipped with a DMA-RH environmental control accessory. Frequency sweeps (0.1 to 100 Hz) were recorded at 55%, and 98% of relative humidity (RH). Tests were conducted in displacement control mode with an oscillation amplitude of 0.001 mm, corresponding to a dynamic strain of approximately *∼*0.02%

### 2.3 Data analysis and modeling

#### 2.3.1 Dynamic properties and Surrogate Modeling

Vibration response data were analyzed using MATLAB’s tfestimate function within a custom-developed script to determine transfer functions correlating input and output signals. Dynamic mechanical properties, specifically the dynamic modulus *E*_d_ and loss factor *η*, were calculated from frequency response data from the following relationships [2]:

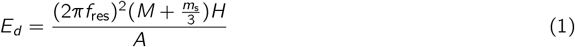

In 1, *f*_res_ is the resonance frequency, *m*_s_ is the hydrogel’s mass, *M* is the top mass, *H* is the thickness and *A* is the cross-sectional area of the gel samples. The loss factor *η* was calculated as:

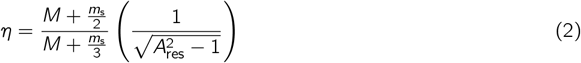

The term *A*_res_ in 2 represents the amplitude at the resonance frequency. The loss modulus was subsequently determined as the product of the dynamic modulus and the loss factor [17].

#### 2.3.2 Compressive properties and hyperelastic modeling

The Young’s modulus (*E*) was determined from the linear region of the stress–strain curve, specifically near the origin (i.e., for strains < 0.05)[33, 34].

To describe the large-strain behavior, the compression data were fitted to a two-parameter Mooney–Rivlin model of hyperelasticity [35]. The relationship between the uniaxial stress (*σ*), the material parameters (*C*_01_ and *C*_10_), and the elongation (*λ*) is defined as:

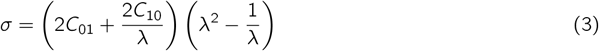

The material constants *C*_01_ and *C*_10_ were identified from the experimental quasi-static compression data via nonlinear least-square fitting into Equation 3.[35]

To quantify the energy absorption capacity within the compression range, a Modulus of Resilience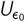 was calculated as the area under the linear stress-strain (*σ − ϵ*) curve up to a fixed strain reference (*ϵ* = *ϵ*_0_):[36, 37]

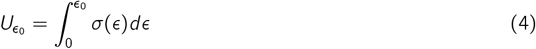

### 2.4 Gaussian Process Regression and model validation

#### 2.4.1 Model training and Response Surfaces

The multidimensional relationship between gel composition (agarose concentration), environmental conditions (relative humidity), loading (top mass), and the resulting viscoelastic properties (*E*^*′*^, *E*^*′′*^, and tan *δ*) was modeled using Gaussian Process Regression (GPR) with Automatic Relevance Determination (ARD) `matérn` 5/2 kernel and a constant basis function after previews standardization.[38, 39]. Although other kernels provide strong numerical fits, the `matérn` 5/2 kernel was selected for its differentiable nature, which captures the smooth, continuous physical transitions intrinsic to the viscoelasticity of agarose gels. Additionally, a constant basis function was used to maintain a strictly non-parametric approach. The final response surfaces and contour plots were generated by training the GPR models on the complete experimental dataset.

#### 2.4.2 Grouped cross-validation procedure

To rigorously assess the predictive accuracy and applicability of the GPR framework—while avoiding overfitting—a grouped *k* -fold cross-validation (*k* = 3) was performed. The dataset was partitioned into three subsets using unique sample identifiers, ensuring all replicates of a given sample remained in the same fold. This “grouped” approach tests the model’s ability to predict properties of entirely new gel formulations, rather than merely interpolating between identical replicates.[40]

#### 2.4.3 Performance metrics and reliability

The mathematical reliability of the generated surfaces has been quantified by the cross-validation Coeffi-cient of Determination 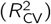. This metric is calculated as:[36]

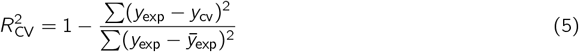

in 5, *y*_exp_ are the experimental values and *y*_*cv*_ are the “blind” predictions generated during the cross-validation cycles.

### 2.5 SEM images

Morphological analyses were performed using a FEI environmental scanning electron microscope (ESEM) equipped with a field emission gun (FEG) at the University of Parma. Images and spectra were recorded at an accelerating voltage of 7.0 kV under high vacuum.

### 2.6 Statistical analysis

Quasi-static compression tests employed five replicates (*n* = 5) to mitigate sensitivity to geometric defects and optimize hyperelastic model fitting.

A one-way analysis of variance (ANOVA), computed via the anova1 function, was performed utilizing the global p-value to confirm the overall statistical significance of the data across all concentrations. DMA and vibration transmissibility were performed using three replicates (*n* = 3) to minimize compositional variability induced by prolonged environmental exposure thereby maintaining experimental consistency.

Consequently, dynamic measurements are reported as mean values accompanied by their minimum and maximum limits. All mathematical modeling and statistical analyses were executed using MATLAB software.[41]

## 3 Results and discussion

### 3.1 Structural architecture and quasi-static compressive properties

The mechanical response of agarose hydrogels is fundamentally linked to their temperature-dependent sol-gel transition. Heating the agarose formulation in an aqueous solution to *∼* 100°*C* induces a highly disordered state of isolated “curly coils” (Fig. 1A). Upon cooling to room temperature, these polymer chains undergo a conformational transition into double helices (Fig. 1B), which subsequently aggregate to form a structurally stable, three-dimensional porous network ((Fig. 1C). Scanning electron microscopy (SEM) of the lyophilized samples (See Supporting Information, Fig. S1) confirms the presence of this architecture, revealing a highly interconnected, porous micro-structure capable of retaining substantial water volumes.

**Figure 1:**
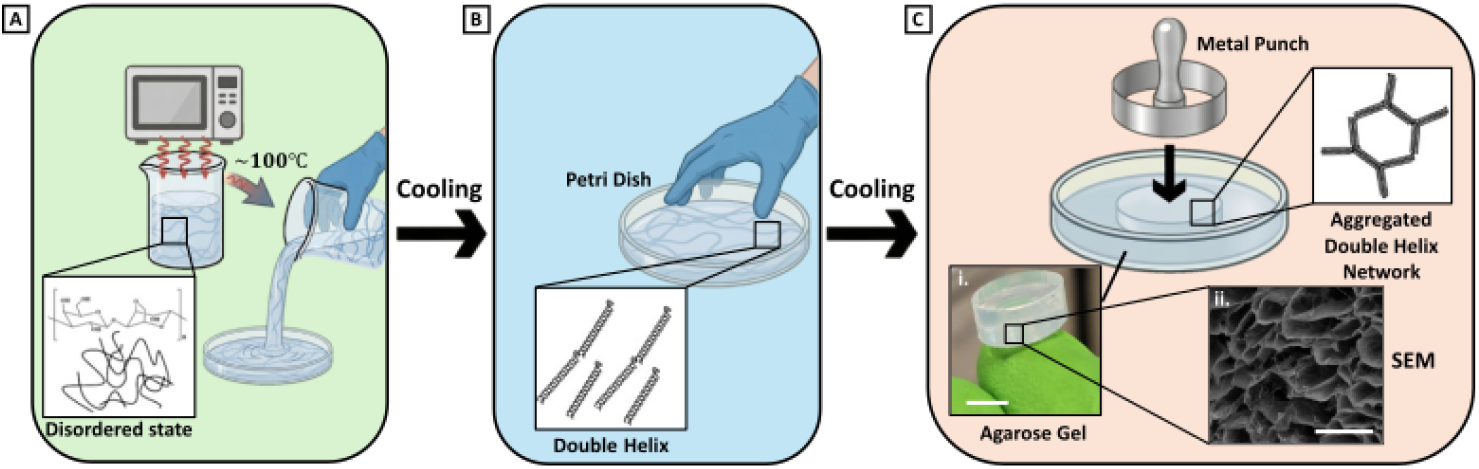
Preparation and structural evolution of agarose hydrogels. **(A)** Heating of agarose powder in aqueous solution to *∼* 100°C via microwave irradiation, resulting in a disordered state. **(B)** Initial cooling phase within a Petri dish, where polymer chains transition into a double-helix conformation. **(C)** Formation of a stable agarose gel through further cooling and the development of an aggregated double-helix network. The final gel is processed using a metal punch for mechanical testing. Lyophilized agaroze is characterized by Scanning Electron Microscopy (SEM). The scale bar represents (i) 100 *µm* for SEM and (ii) 13 mm for the agarose gel at 1%.

The hydrogels were initially evaluated under fully hydrated conditions (98% RH), considered as a baseline configuration for reference properties. Quasi-static uniaxial compression testing demonstrated a predictable, monotonic strain-stiffening effect associated with increasing polymer density. The Young’s modulus, derived from the linear elastic region (5-10% strain, Fig.2A), increased by over an order of magnitude, scaling from 48 *±* 3 kPa at 1 wt% concentration to 1338 *±* 336 kPa at 7 wt% (Fig.1C) [33, 34]). This concentration-dependent stiffening is accompanied by a proportional increase in the modulus of resilience, rising from *∼*1 kPa at 1 wt% to nearly 25 kPa at 7 wt%. This increase is an indication of the network’s enhanced capacity for elastic energy storage under high-hydration states (Fig. 2D).

**Figure 2:**
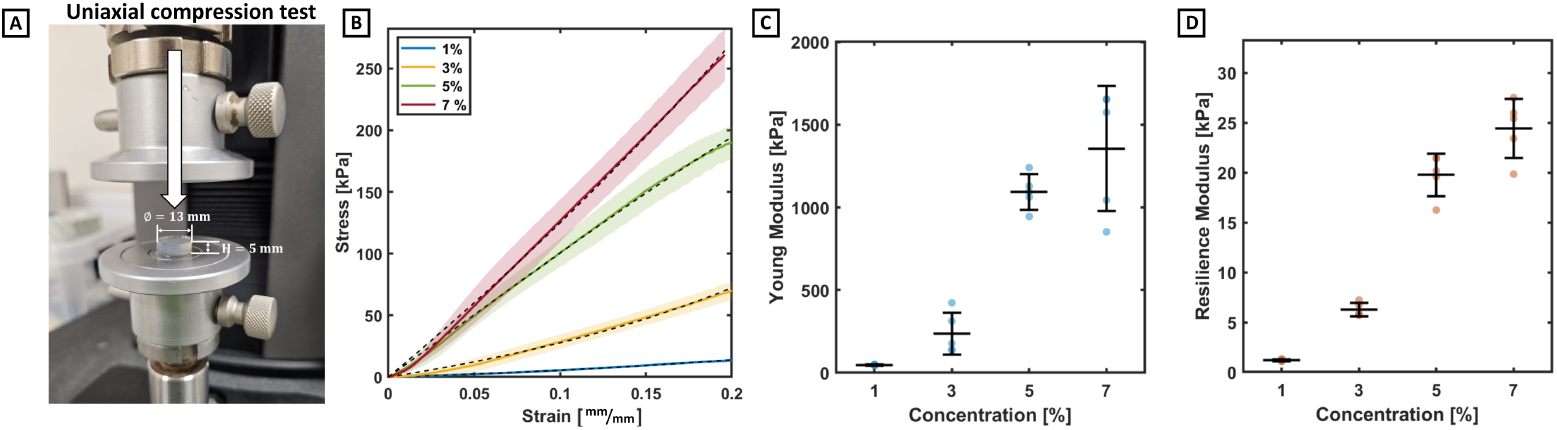
Compression and Mooney-Rivlin fitting of porous agarose hydrogels. (A) Schematic of the uniaxial compression test. The indicated height of 5 mm is a representative nominal value; actual specimen dimensions were recorded for each test to calculate stress and strain (see Supporting information, Tab. S2). (B) Average stress-strain curves for hydrogels at varying agarose concentrations (1%, 3%, 5%, and 7%), obtained from *n* = 5 independent replicates. Solid lines represent the mean data, while shaded areas indicate the standard deviation. Dashed lines represent the Mooney-Rivlin fitting applied to the averaged experimental data. All fits show excellent agreement with the data (*R*^2^ *>* 0.99 for 1%, 3%, 5% and 7%). (C) Young’s moduli from compression tests (compression speed 0.6 mm/min).(D) Modulus of resilience calculated at *ε*_*max*_ = 0.20 for different agarose concentrations. Data in (C) and (D) are presented as individual data points (scatter) with mean horizontal lines; error bars denote the standard deviation. Statistical significance across groups in (C) and (D) was evaluated using a One-Way ANOVA (global *p <* 0.01 indicates a significant effect of concentration).

**Figure 3:**
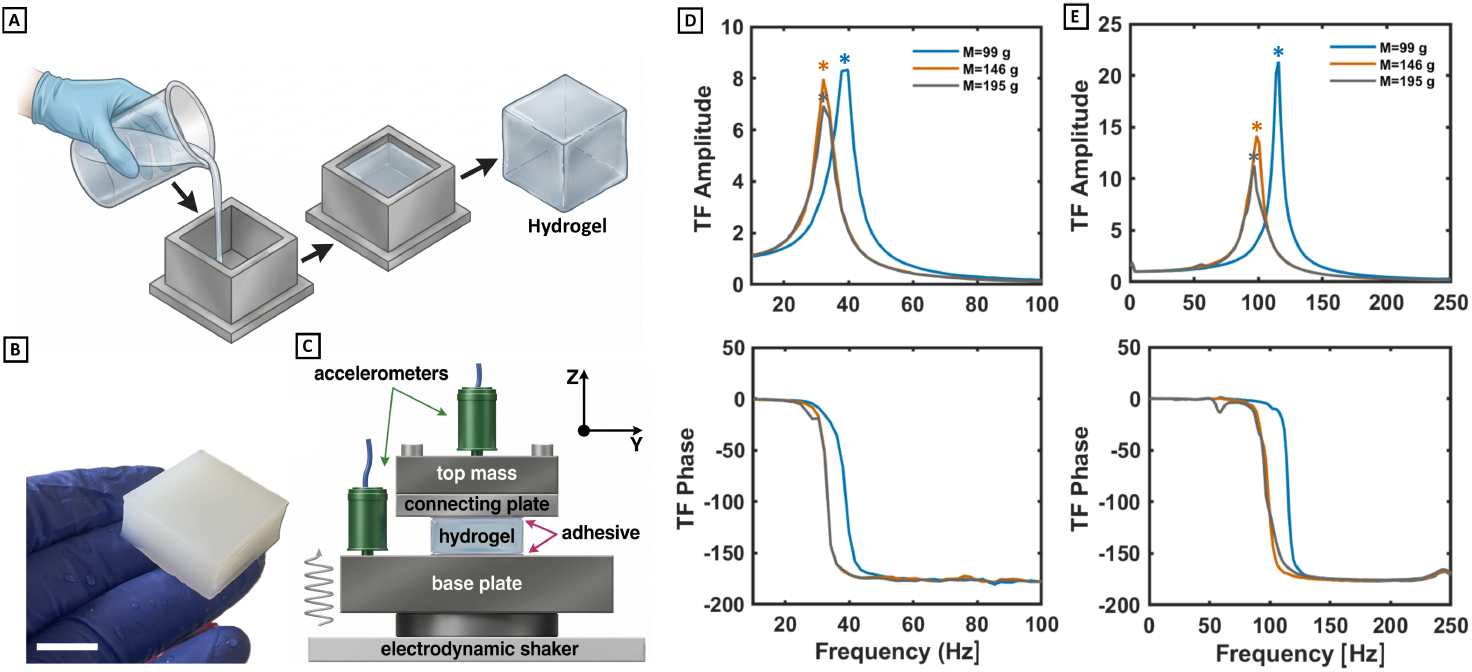
Vibrational transmissibility properties of agarose hydrogels at different concentrations. **(A)** Hydrogel block for vibration transmissibility experiments were prepared using a mold. **(B)** Optical image of a hydrogel block of dimensions 30 *×* 30 *×* 15 mm, scale bar demarcates 15 mm. **(C)** Schematic of the experimental setup. A hydrogel block is secured between a base plate and a connecting plate using alginate tray adhesive (magenta). To apply incremental compressive prestress, interchangeable top masses are secured to the connecting plate via screws. Accelerometers (green) mounted on the base plate and top mass monitor mechanical wave amplitudes before and after transmission through the hydrogel. **(D-E)** Vibration transmissibility data. Representative transmissibility factor (TF) amplitude curves of agarose 1% and 3% (top row) with the corresponding TF phase curves (bottom row) at 98% of humidity. Asterisks indicate resonant frequency.

### 3.2 Hyperelastic behavior

Elastomeric and gel-like materials used in vibration damping are routinely subjected to large, non-linear strains and quasi-static forces [17]. The experimental data from uniaxial compression were therefore further fitted to a two-parameters Mooney-Rivlin model that describe a baseline hyperelastic behavior. The experimental stress-strain curves shows good agreement with the model across all concentrations, with highly reliable fits (*R*^2^ *>* 0.99). The material parameters (*C*_10_ and *C*_01_) describe the uniaxial large-strains compression behavior of these agarose gels. By applying the relationship *E* = 6(*C*_10_ + *C*_01_), we also derived an expression for the theoretical Young’s moduli (Tab. 2).

**Table 1:** Mechanical and viscoelastic properties of agarose hydrogels. The fully hydrated state (98% RH) is compared to the damping state (75% RH) at a fixed top mass of *M* = 99 g.

| Conc.<br>[%] | 98% RH |  |  |  | 75% RH |  |  |
| --- | --- | --- | --- | --- | --- | --- | --- |
| | $E$ [kPa] | $E'$ [kPa] | $E''$ [kPa] | $\tan \delta$ | $E'$ [kPa] | $E''$ [kPa] | $\tan \delta$ |
| 1 | $47 \pm 3$ | 107 | 13 | 0.12 | 86 | 13 | 0.14 |
| 3 | $227 \pm 135$ | 923 | 44 | 0.05 | 1107 | 235 | 0.21 |
| 5 | $1076 \pm 141$ | 3118 | 243 | 0.08 | 2372 | 515 | 0.21 |
| 7 | $1339 \pm 398$ | 2734 | 173 | 0.06 | 3695 | 283 | 0.08 |

**Table 2:** Mooney-Rivlin parameters for agarose hydrogels. Comparison of *C*_01_ and *C*_10_ coefficients across different concentrations.

| Parameter<br>[kPa] | Concentration [%] |  |  |  |
| --- | --- | --- | --- | --- |
|  | 1 | 3 | 5 | 7 |
| $C_{01}$ | 6.9 | 42.4 | -163.5 | -65.4 |
| $C_{10}$ | 0.6 | -5.7 | 331.8 | 259.4 |
| $E = 6(C_{10} + C_{01})$ | 45 | 220 | 1010 | 1164 |

The consistency between the theoretical predictions and the experimental data confirms the model’s accuracy, with errors of 4.2%, 3.1%, 6.1%, and 13.1% across the 1-7% agarose concentration range. This close alignment confirms that the Mooney-Rivlin framework accurately describes the hyperelasticity and load-bearing capacity of the agarose matrix.

### 3.3 The influence of hydration

Fig. 4 shows the relation between loss and the dynamic moduli at different values of humidity and prestress. While the fully hydrated state (98% RH) emphasizes the network’s elastic storage capacity, effective vibration mitigation depends primarily on viscous energy dissipation. Damping efficiency depends on the balance between network stiffness and polymer chain friction. Our dynamic transmissibility tests—conducted at 55%, 75%, and 98% RH—show this interplay is highly sensitive to environmental conditions. At full saturation (98% RH), the dynamic modulus (*E*^*′*^) reaches up to *∼*3 MPa at 7 wt% concentration, but the water heavily lubricates the polymer chains, restricting viscous friction. Consequently, the loss factor (tan(*δ*)) is surprisingly low, dropping from 0.12 at 1 wt% to a minimal 0.06 at 7 wt %. To validate these trends, DMA compressive tests were conducted at 28 Hz (see Supporting Information, Tab. S10, S11) — matching the fundamental resonance of the 1.0 wt% system — while avoiding parasitic and inertial resonances in the measurement setup. This cross-validation was restricted to the 1.0 wt% formulation to ensure clear and unambiguous material characterization. DMA tests of 1.0 wt % agarose at 98% of humidity exhibit a storage modulus of 32 *±*7 kPa close to the value of Young’s modulus (47 *±* 3 kPa) while, the vibration transmissibility tests under a 99 g top mass yielded a significantly larger dynamic stiffness (*E*^*′*^ *≈* 107 kPa). However, this divergence arises from the static prestress applied to the material due to the presence of the top mass. The tangent modulus from quasi-static compression curves at the corresponding applied stress of 7.3 kPa (corresponding to the 99 g top mass) yields a value of 79.5 kPa for the 1.0 wt% gel (see Supporting information, Tab. S3). Dynamic stiffening up to 7-10 times — where compressive properties vary with frequencies below 300 Hz during vibration transmissibility tests—has been previously reported in hydrogel systems such as alginate/poloxamer[2, 16]. In this case, the dynamic modulus at a very low frequency (28 Hz) shows a *∼*34% increase to the quasi-static modulus. The substantial alignment of the tangent modulus with the storage modulus *E*^*′*^ reflects the non-linear response of the gel to the applied static load. A comprehensive comparison of these modulus ratios across all concentrations at 98% RH is summarized in Table S2 of the Supporting Information. Conditioning at an intermediate 75% RH produces a highly non-monotonic damping response, dramatically amplifying viscous dissipation [42]. Both 3 and 5 wt% gels reach a peak tan *δ* of 0.21, with the 5 wt% formulation maximizing the loss modulus (*E*^*′′*^ = 514.9 kPa, more than double its fully hydrated state). At this intermediate hydration level, internal chain friction is optimized while molecular rearrangement remains active, achieving damping efficiency comparable to synthetic elastomers [30, 43]. However, dehydration to 55% RH significantly restricts chain mobility, resulting in pronounced network stiffening. The 7 wt% *E*^*′*^ exceeds 5 MPa under high pre-stress, while structural locking causes the previously optimal 5 wt% gel’s *E*^*′′*^ to decrease to *∼*200 kPa. At 55% RH, only the densest 7 wt% network dissipates substantial energy (*∼*450 kPa), behaving predominantly as a glassy solid. Mechanistically, this behavior suggests that environmental conditioning drives the polymer network across a rugged chemical potential landscape. In the solvent-rich regime (98% RH), an excess of entropic degrees of freedom leads to extensive plasticization; here, the aqueous phase effectively screens interchain forces, suppressing damping efficiency. Conversely, at low humidity (55% RH), the system compensates for reduced configurational entropy through robust enthalpic interactions, which constrain the network into a kinetically trapped state. The dissipation peak observed at 75% RH thus emerges as a critical transition point near the global potential minimum. In this intermediate regime, humidity-induced local ordering optimizes internal friction by balancing chain mobility with intermolecular constraints, bypassing both the plasticization limit and enthalpic arrest.[44] Thus, optimal vibration damping is achieved by targeting the 75% RH regime—not by simply increasing polymer concentration. Comprehensive loss modulus values across the investigated pre-stress levels, concentrations, and humidity conditions are detailed in Table S7 of the Supporting Information.

**Figure 4:**
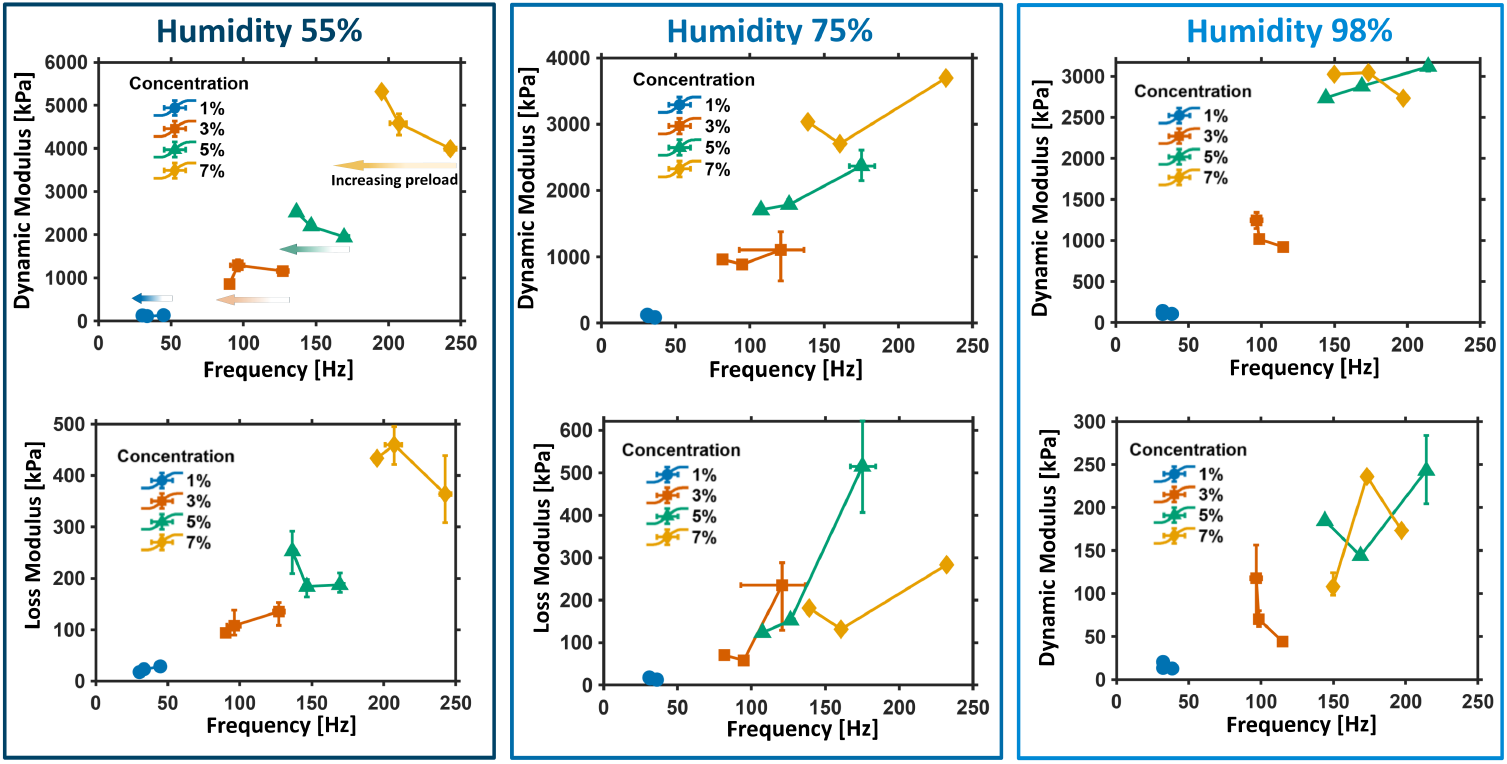
Viscoelastic characterization of agarose hydrogels under different environmental and mechanical conditions. Dynamic mechanical analysis of agarose hydrogels across a concentration range of 1-7% (wt %) and relative humidity levels (55, 75 and 98%). The storage modulus and the loss modulus are plotted as a function of resonance frequency. From left to right, the three columns represent increasing humidity levels. In each plot, for a given concentration curve, movement from right to left indicates increasing applied top mass (i.e., rising prestress) respectively of 99 g, 146 g and 195 g. Error bars represent the minimum and maximum measured values for *n* = 3 independent samples.

### 3.4 Mapping the dynamic properties via predictive modeling

The Gaussian Process Regression effectively modeled the complex interactions observed in the vibration transmissibility test data by providing accurate predictive surfaces (grouped cross-validated 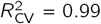 for dynamic modulus and 0.92 for loss modulus; see Table S8 for the complete evaluation of GPR architectures). It is important to note that these predictive surfaces do not map the viscoelastic response at a single fixed frequency; rather, they represent the equivalent dynamic properties evaluated at the fundamental resonance frequency specific to each hydrogel configuration and applied pre-load (the GPR predictive mapping, uncertainty analysis, and model validation for resonance frequencies are provided in Figs. S4, S5 and Table S9).

The contour maps (Fig. 5) reveal a qualitative trend in dynamic stiffness: it peaks at 7 wt% concentration and 55% RH, reaching values up to 5 MPa under high prestress. To assess the statistical reliability of these trends, the associated predictive uncertainty maps and 3D response surfaces are reported in Figs. S2 and S3, respectively. In contrast, the behavior of the energy absorbed by damping (i.e., the equivalent loss modulus here) is more complex. Energy dissipation does not increase steadily; instead, it peaks in specific locations optimizing at 3–5 wt% concentration and 75% RH under low prestress. The GPR model shows that the damping changes with the load. As static prestress increases, the optimal damping surface peaks shift to lower polymer concentrations (1–3 wt%). This suggests that heavy static loads over-compress denser networks, thereby limiting internal chain friction. As a consequence, the optimal energy dissipation belongs to softer, less concentrated gel architectures.

**Figure 5:**
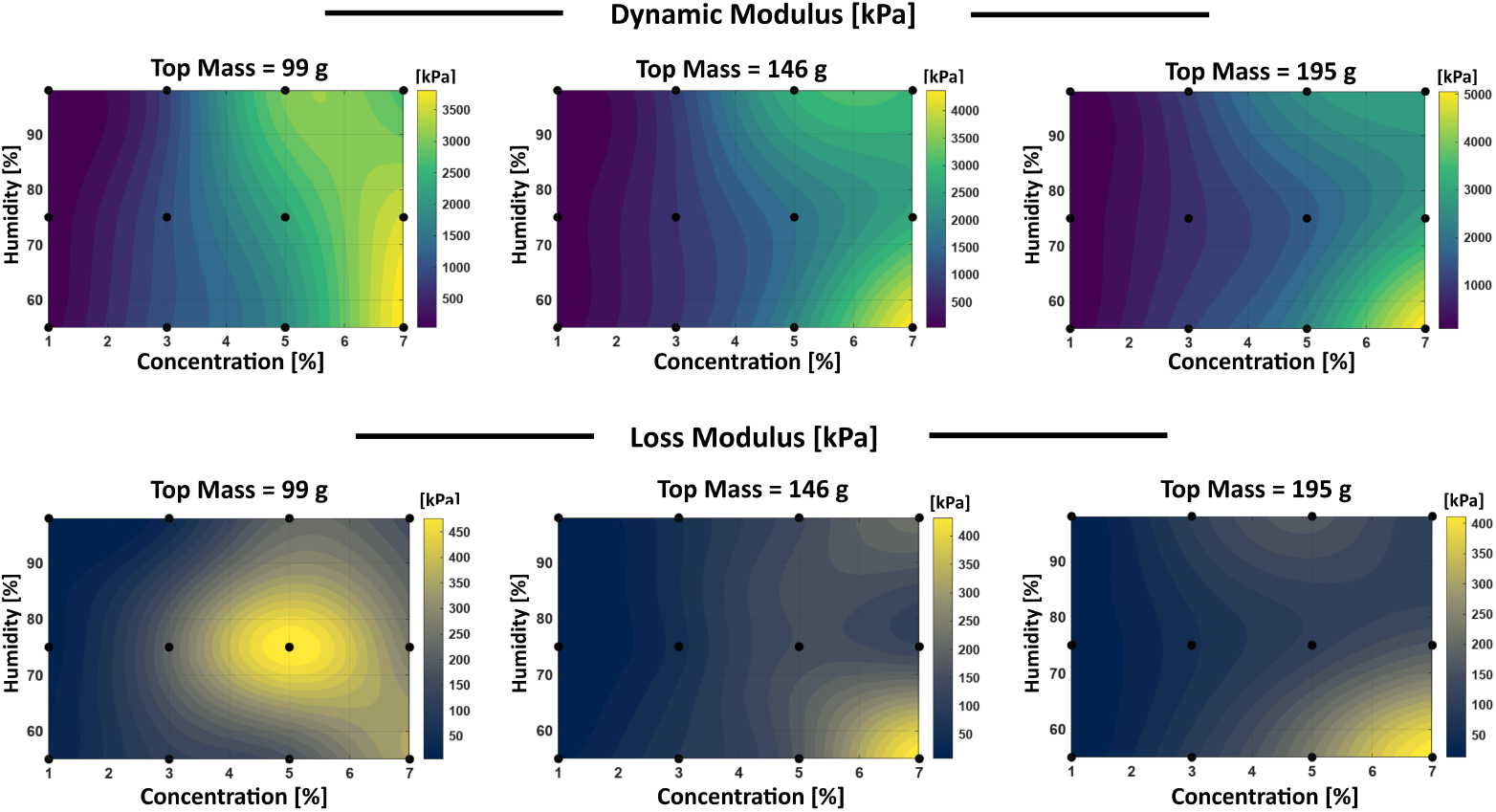
Gaussian Process Regression (GPR) predictive surfaces of hydrogel viscoelasticity. Contour maps illustrating the predicted Dynamic Modulus (top row, Grouped CV RMSE: 122 kPa and Grouped 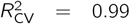, and Loss Modulus (bottom row, Grouped CV RMSE: 37 kPa and Grouped CV 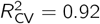of agarose hydrogels as a continuous function of polymer concentration and relative humidity. The columns, from left to right, represent incrementally increasing compressive pre-stress (top mass) applied to the samples. Black markers indicate the specific experimental data coordinates used to train the GPR model. The model uses an Automatic Relevance Determination (ARD) `matérn` 5/2 kernel to capture the highly nonlinear, multidimensional dependency between mechanical damping and dynamic stiffness properties on hydration, network density and static pre-load.

To obtain further insights into the behavior of the dynamic properties of these agarose gels, the GPR surfaces were differentiated and partial derivatives were calculated numerically via the finite difference method using the MATLAB built-ingradient function to yield sensitivity maps at a constant 99 g of mass preload (Fig. 6).[41] Viscoelastic sensitivity maps evaluated under the full range of pre-stress masses, from 99 g to 195 g, are reported in Fig. S6 of the Supporting Information. Analyzing partial derivatives with respect to polymer concentration (*∂/∂C*, top row) and relative humidity (*∂/∂RH*, bottom row) converts property maps into a stability analysis of the hydrogel networks under these dynamic conditions.

**Figure 6:**
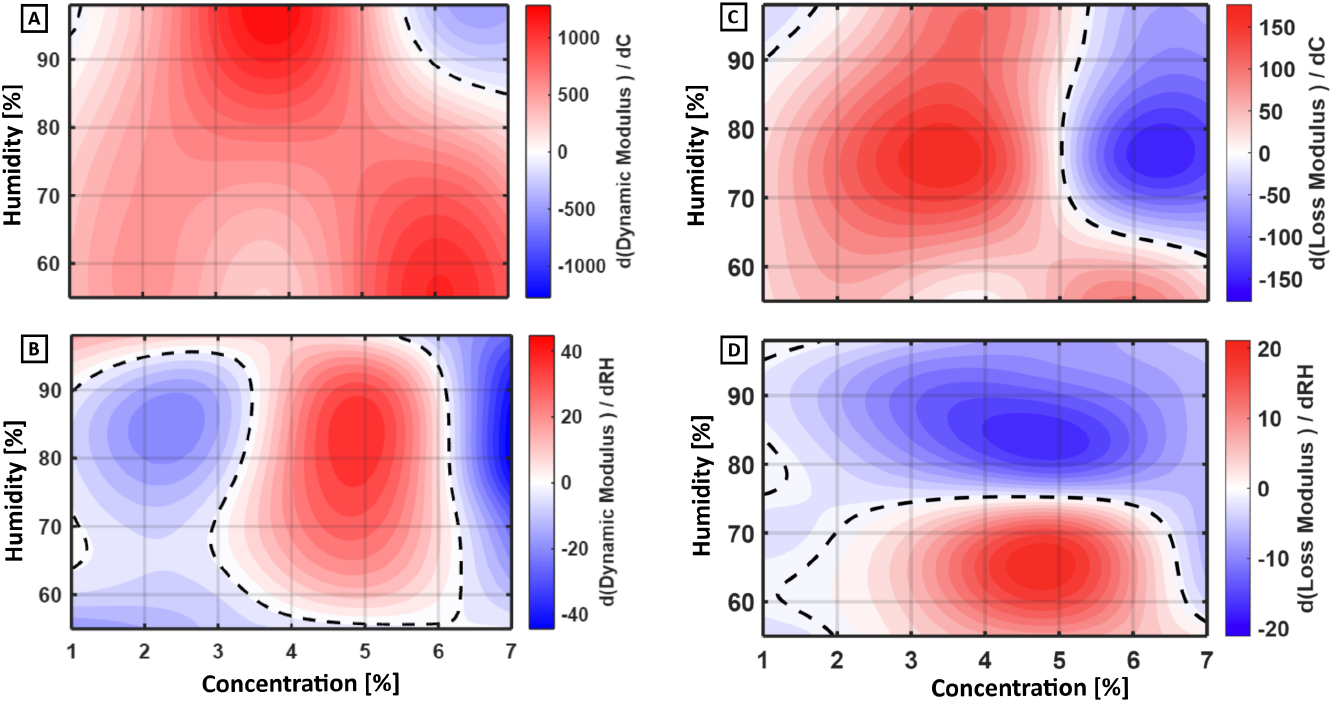
Viscoelastic sensitivity maps of agarose hydrogels evaluated under a constant preload mass of 99 g. The contour plots visualize the partial derivatives of the material’s dynamic response with respect to polymer concentration (top row: A, C) and relative humidity (bottom row: B, D). Columns correspond to the Dynamic Modulus (A, B), Loss Modulus (C, D). Divergent color scales indicate the local rate of change, with red and blue regions denoting positive and negative gradients, respectively. Dashed black lines represent zero-gradient contours, mathematically delineating critical structural and environmental transition zones where the material properties reach a local maximum, minimum, or plateau.

Here, zero-gradient contours (dashed lines) define critical transition zones where damping efficiency remains robust against environmental changes. By contrast, steep positive (red) or negative (blue) gradients indicate regions where minor environmental changes induce rapid mechanical stiffening or a sudden collapse in internal chain friction.

The dynamic modulus reveals distinct stiffness sensitivities. Increasing polymer concentration globally stiffens the network (Fig. 6A), particularly at hight levels of hydration. However, the network’s moisture response (Fig. 6B) is highly nonlinear. In leaner networks, increased humidity acts as a bulk lubricant, rapidly softening the gel. At intermediate concentrations (4–5 wt%), a slight initial stiffening precedes ultimate softening. The loss modulus maps define boundaries of optimal damping efficiency. Fig. 6C identifies a stability plateau at intermediate concentrations, bordered by a steep negative gradient (>5 wt%) where excessive polymer density rigidly locks the internal chains, decreasing damping capability. The figure also indicates the fundamental role of hydration, with a sharp transition at *∼*75% RH (Fig. 6D). Below this threshold, added moisture liberates restricted chains, enhancing damping. Beyond it, the network becomes over-lubricated, causing damping efficiency to collapse. These results confirm that while drier gels need moisture to enable inter-chain friction, oversaturation inhibit the polymer-polymer interactions essential for effective energy dissipation.

## Conclusions

This study reveals that the dynamic mechanical behavior of agarose hydrogels under varying environmental conditions is not solely determined by polymer concentration. Instead, it is critically influenced by the interplay of network density, environmental hydration, and mechanical prestress. While the hydrogels’ network stiffness increases proportionally with the polymer fraction, damping exhibits a highly nonlinear dependency on environmental conditioning. We identified a structural optimal state at intermediate hydration (75% RH) and network density (3 wt%), where the balance between polymer chain mobility and internal friction maximizes energy dissipation, yielding a loss factor (tan(*δ*)) of 0.21 at the resonance frequency of 175 Hz – performance directly comparable to conventional synthetic elastomers. Gaussian Process Regression (GPR) revealed that the mapping of the dynamic properties is load-dependent. Increased mechanical prestress reshapes the damping landscape, shifting the optimal energy dissipation state from denser networks toward leaner, more compliant polymer architectures (as low as 1 wt%).

These findings demonstrate that agarose hydrogels can be highly adaptive viscoelastic systems. By systematically mapping the influence of hydration and static preload on the internal hydrogels’ network friction, this work establish mechanistic design principles for engineering sustainable, high-performance soft materials tailored for vibration alleviation.

## Author contributions

**Ikenna Ojoboh**: Methodology, Investigation, Formal analysis, Writing - original draft. **Manuel Dedola**: Investigation, Formal analysis, Data curation, Validation, Writing - original draft. **Katherine Nelms**: Methodology, Supervision, Validation, Project Administration. **Charles de Kergariou**: Methodology, Formal analysis, Data curation, Resources. **Ibrahim Patrick**: Methodology, Supervision. **Ludovico Cademartiri**: Supervision, Formal Analysis. **James P. K. Armstrong**: Supervision, Resources. **Adam W. Perriman**: Conceptualization, Funding Acquisition. **Fabrizio Scarpa**: Conceptualization, Supervision, Resources, Project administration, Funding acquisition, Writing - review and editing.

## Data availability

The data supporting the findings of this study are available within the article and its supporting information. Raw data are available from the corresponding author upon reasonable request.

## Acknowledgements

KN, MD, CdK, FS and AWP acknowledge the support from the H2020 ERC-2020-AdG 101020715 NEUROMETA project. CdK also acknowledges the EPSRC Doctoral Prize Fellowship for supporting this work (Grant No. EP/W524414/1). MD and LC acknowledge funding from the Associazione Italiana di Cristallografia (AIC) through the ‘Fiorenzo Mazzi’ Scholarship. JPKA acknowledges funding from a UKRI Future Leaders Fellowship (MR/V024965/1)

## S1 SEM OF AGAROSE SAMPLES

**Figure S1:**
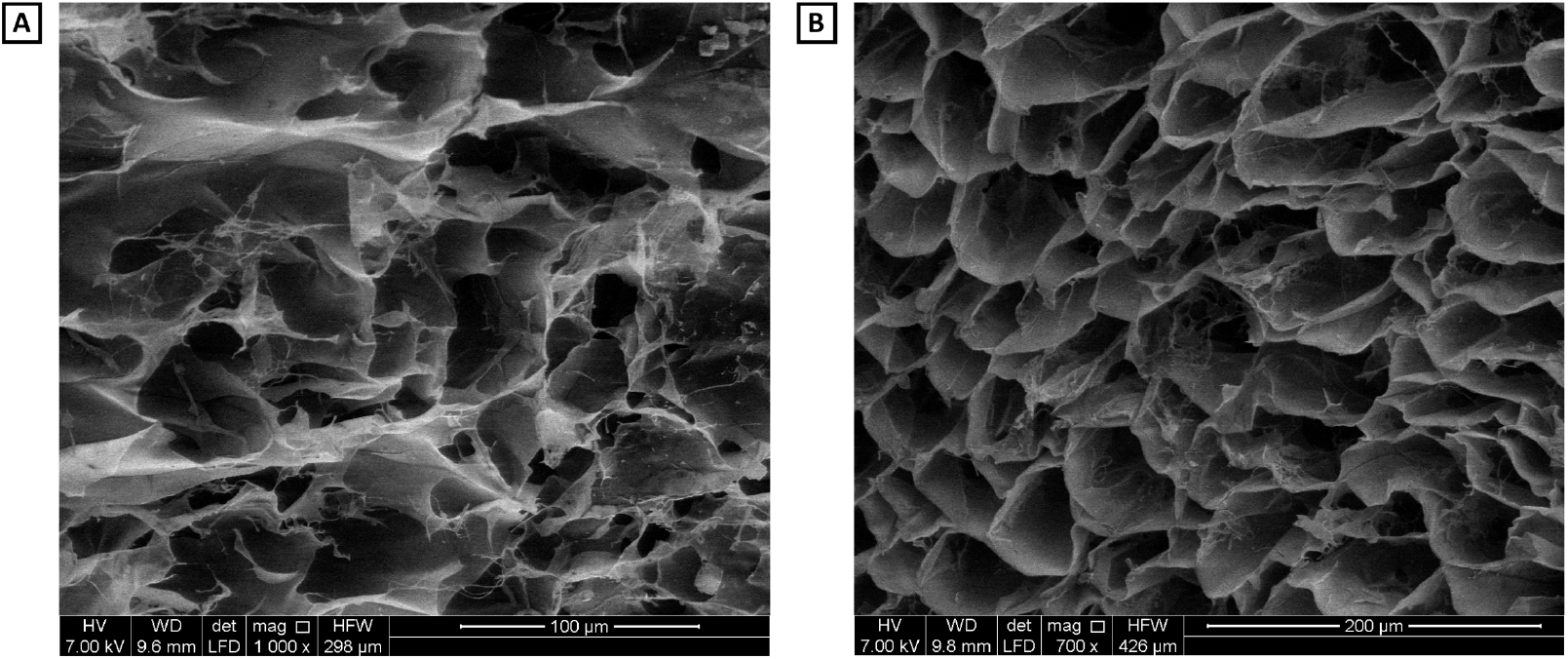
Scanning electron micrographs (SEM) of lyophilized 1 wt% agarose. **(A)** Surface morphology. **(B)** Internal cross-section of the sample.

## S2 SUMMARY OF EXPERIMENTAL MODULUS COMPARISONS

**Table S1:**
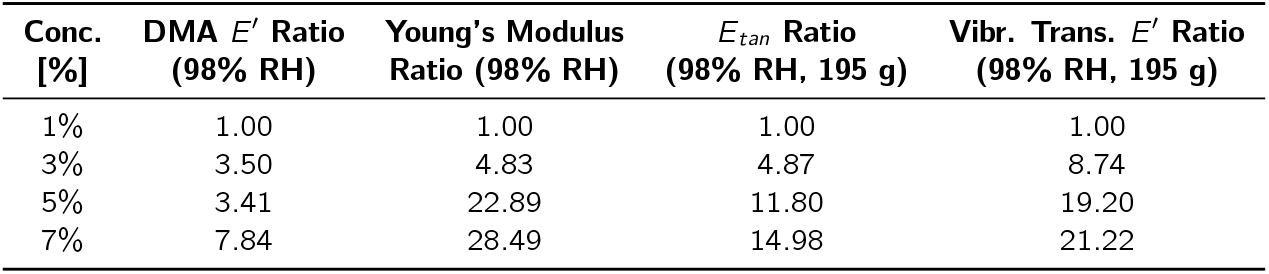
Summary of modulus ratios at 98% RH (*E*(*x* %)*/E*(1%), comparing DMA storage modulus (*E*^*′*^), Young’s modulus, tangent modulus (*E*_*tan*_), and vibration transmissibility dynamic modulus (*E*^*′*^) under preload mass of 195 g.

## S3 AGAROSE SAMPLE DATA: QUASISTATIC COMPRESSION TESTS

**Table S2:**
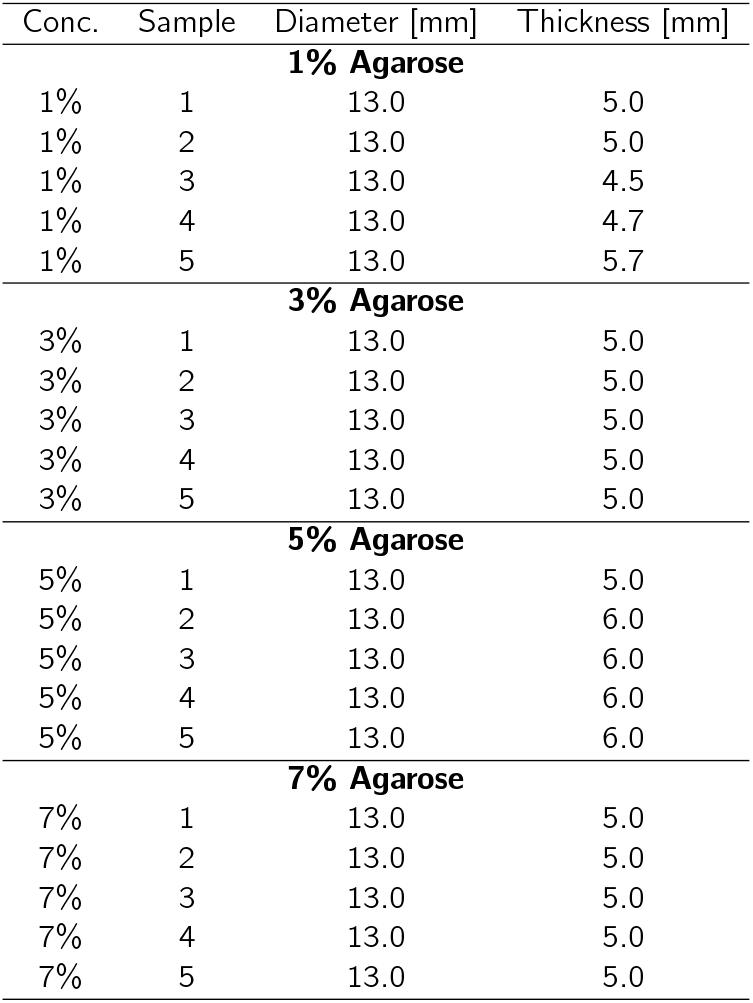
Thickness [mm] used in Quasistatic compression (compression speed 0.6 mm/min).

| Conc. | Sample | Diameter [mm] | Thickness [mm] |
| --- | --- | --- | --- |
| **1% Agarose** |  |  |  |
| 1% | 1 | 13.0 | 5.0 |
| 1% | 2 | 13.0 | 5.0 |
| 1% | 3 | 13.0 | 4.5 |
| 1% | 4 | 13.0 | 4.7 |
| 1% | 5 | 13.0 | 5.7 |
| **3% Agarose** |  |  |  |
| 3% | 1 | 13.0 | 5.0 |
| 3% | 2 | 13.0 | 5.0 |
| 3% | 3 | 13.0 | 5.0 |
| 3% | 4 | 13.0 | 5.0 |
| 3% | 5 | 13.0 | 5.0 |
| **5% Agarose** |  |  |  |
| 5% | 1 | 13.0 | 5.0 |
| 5% | 2 | 13.0 | 6.0 |
| 5% | 3 | 13.0 | 6.0 |
| 5% | 4 | 13.0 | 6.0 |
| 5% | 5 | 13.0 | 6.0 |
| **7% Agarose** |  |  |  |
| 7% | 1 | 13.0 | 5.0 |
| 7% | 2 | 13.0 | 5.0 |
| 7% | 3 | 13.0 | 5.0 |
| 7% | 4 | 13.0 | 5.0 |
| 7% | 5 | 13.0 | 5.0 |

**Table S3:** Tangent modulus under varying pre-stress conditions, calculated from mean stress-strain curves.

| Conc. | Mass [kg] | Stress [kPa] | Strain [mm/mm] | Tangent Modulus [kPa] |
| --- | --- | --- | --- | --- |
| 1% | 0.099 | 7.3 | 0.124 | 79 |
| 1% | 0.146 | 10.8 | 0.164 | 89 |
| 1% | 0.195 | 14.4 | 0.211 | 82 |
| 3% | 0.099 | 7.3 | 0.041 | 288 |
| 3% | 0.146 | 10.8 | 0.052 | 334 |
| 3% | 0.195 | 14.4 | 0.062 | 399 |
| 5% | 0.099 | 7.3 | 0.010 | 806 |
| 5% | 0.146 | 10.8 | 0.014 | 897 |
| 5% | 0.195 | 14.4 | 0.018 | 968 |
| 7% | 0.099 | 7.3 | 0.012 | 924 |
| 7% | 0.146 | 10.8 | 0.015 | 1138 |
| 7% | 0.195 | 14.4 | 0.018 | 1228 |

## S4 AGAROSE SAMPLES DATA: VIBRATION TRANSMISSIBILITY TESTING

**Table S4:**
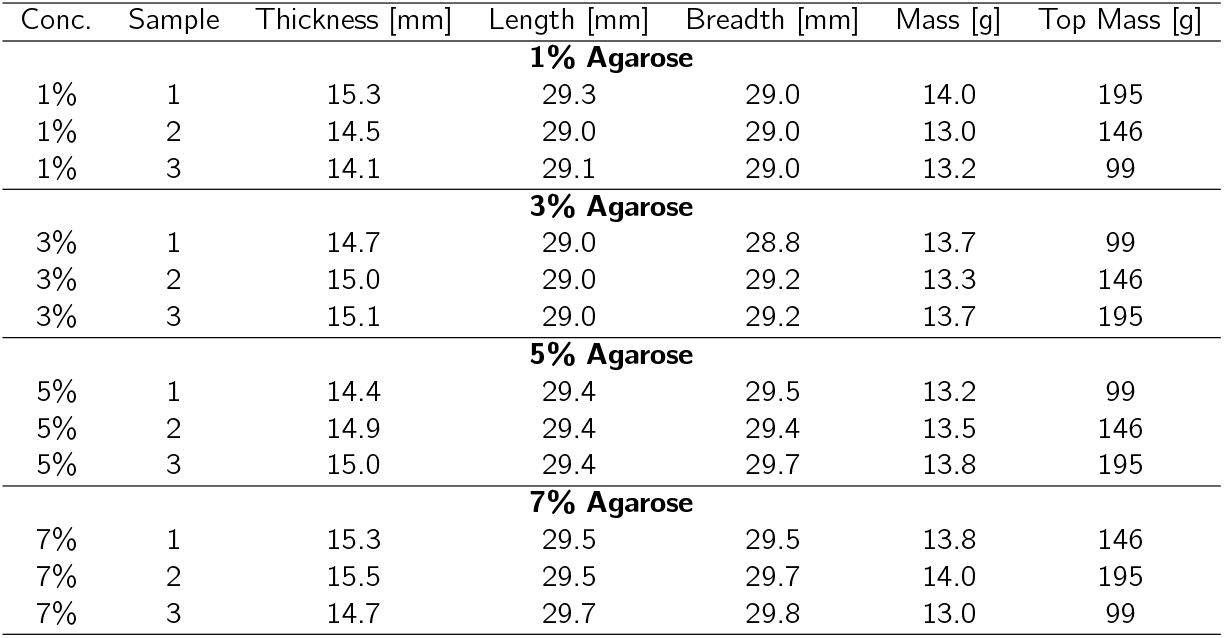
Agarose Samples data at 20°C and 55% RH.

| Conc. | Sample | Thickness [mm] | Length [mm] | Breadth [mm] | Mass [g] | Top Mass [g] |
| --- | --- | --- | --- | --- | --- | --- |
| **1% Agarose** |  |  |  |  |  |  |
| 1% | 1 | 15.3 | 29.3 | 29.0 | 14.0 | 195 |
| 1% | 2 | 14.5 | 29.0 | 29.0 | 13.0 | 146 |
| 1% | 3 | 14.1 | 29.1 | 29.0 | 13.2 | 99 |
| **3% Agarose** |  |  |  |  |  |  |
| 3% | 1 | 14.7 | 29.0 | 28.8 | 13.7 | 99 |
| 3% | 2 | 15.0 | 29.0 | 29.2 | 13.3 | 146 |
| 3% | 3 | 15.1 | 29.0 | 29.2 | 13.7 | 195 |
| **5% Agarose** |  |  |  |  |  |  |
| 5% | 1 | 14.4 | 29.4 | 29.5 | 13.2 | 99 |
| 5% | 2 | 14.9 | 29.4 | 29.4 | 13.5 | 146 |
| 5% | 3 | 15.0 | 29.4 | 29.7 | 13.8 | 195 |
| **7% Agarose** |  |  |  |  |  |  |
| 7% | 1 | 15.3 | 29.5 | 29.5 | 13.8 | 146 |
| 7% | 2 | 15.5 | 29.5 | 29.7 | 14.0 | 195 |
| 7% | 3 | 14.7 | 29.7 | 29.8 | 13.0 | 99 |

**Table S5:**
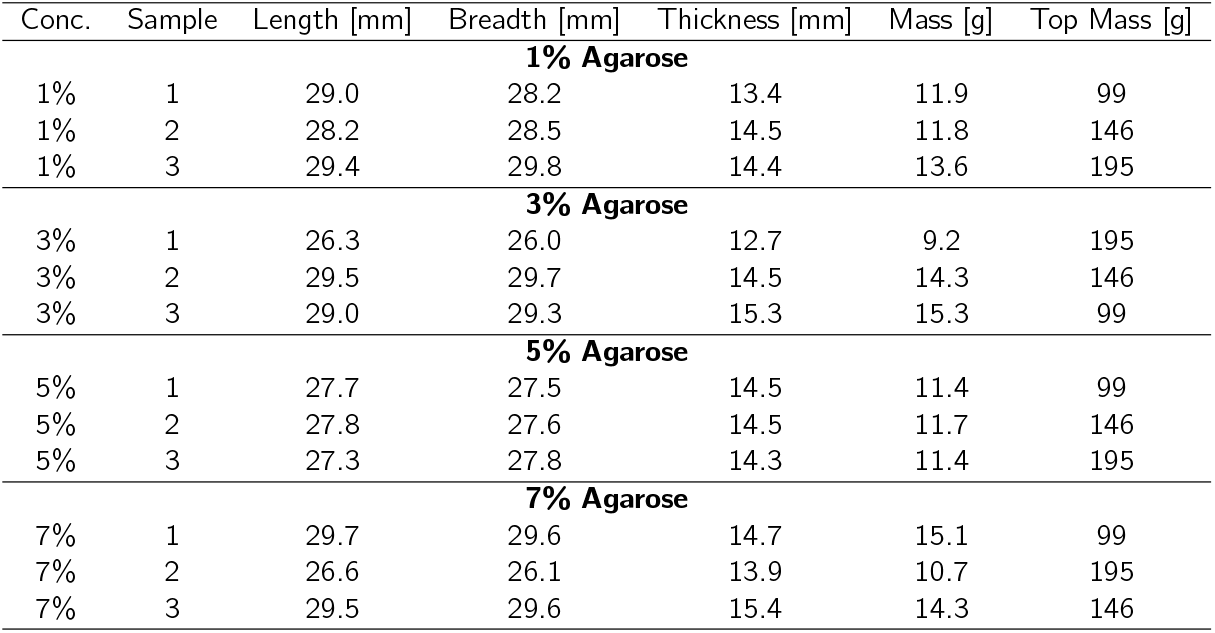
Agarose Samples data at 20.7°C, Sample RH = 75%.

**Table S6:** Agarose Samples data at 20.2°C, Sample RH = 98%.

| Conc. | Sample | Length [mm] | Breadth [mm] | Thickness [mm] | Mass [g] | Top Mass [g] |
| --- | --- | --- | --- | --- | --- | --- |
| **1% Agarose** |  |  |  |  |  |  |
| 1% | 1 | 29.0 | 28.7 | 14.7 | 13.3 | 99 |
| 1% | 2 | 29.0 | 29.1 | 14.6 | 13.4 | 146 |
| 1% | 3 | 29.1 | 28.8 | 14.6 | 13.3 | 195 |
| **3% Agarose** |  |  |  |  |  |  |
| 3% | 1 | 29.5 | 29.6 | 14.9 | 13.5 | 99 |
| 3% | 2 | 29.2 | 29.4 | 15.1 | 13.6 | 146 |
| 3% | 3 | 29.5 | 29.5 | 14.7 | 13.3 | 195 |
| **5% Agarose** |  |  |  |  |  |  |
| 5% | 1 | 29.7 | 29.6 | 14.6 | 13.7 | 99 |
| 5% | 2 | 29.7 | 29.7 | 15.0 | 13.8 | 146 |
| 5% | 3 | 29.7 | 29.6 | 14.7 | 13.7 | 195 |
| **7% Agarose** |  |  |  |  |  |  |
| 7% | 1 | 29.2 | 29.2 | 14.7 | 12.9 | 99 |
| 7% | 2 | 29.1 | 29.3 | 14.6 | 12.9 | 146 |
| 7% | 3 | 29.4 | 29.4 | 14.8 | 13.1 | 195 |

**Table S7:** Mean Loss Modulus [kPa] of agarose hydrogels as a function of concentration and relative humidity for different top masses. Data are presented as Mean (Min - Max) for *n* = 3.

| Top Mass [g] | Conc. [%] | RH 55% | RH 75% | RH 98% |
| --- | --- | --- | --- | --- |
| 99 | 1 | 29 (22 - 43) | 13 (12 - 13) | 13 (13 - 13) |
|  | 3 | 135 (116 - 162) | 235 (125 - 443) | 44 (42 - 46) |
|  | 5 | 187 (173 - 211) | 515 (407 - 621) | 243 (205 - 284) |
|  | 7 | 364 (211 - 568) | 283 (251 - 309) | 173 (17 - 179) |
| 146 | 1 | 23 (19 - 29) | 15.8 (15 - 16) | 14 (13 - 14) |
|  | 3 | 94 (91 - 97) | 58 (55 - 61) | 70 (68 - 71) |
|  | 5 | 184 (165 - 198) | 152 (146 - 161) | 144 (144 - 146) |
|  | 7 | 459 (420 - 489) | 132 (131 - 133) | 236 (226 - 242) |
| 195 | 1 | 18 (14 - 20) | 18 (17 - 18) | 18 (17 - 19) |
|  | 3 | 108 (107 - 110) | 70 (69 - 71) | 118.4 (111 - 123) |
|  | 5 | 253 (231 - 276) | 123.4 (115 - 130) | 185 (184 - 187) |
|  | 7 | 434 (412 - 470) | 181 (180 - 182) | 108 (103 - 113) |

## S5 GAUSSIAN PROCESS REGRESSION AND MODEL VALIDATION

**Figure S2:**
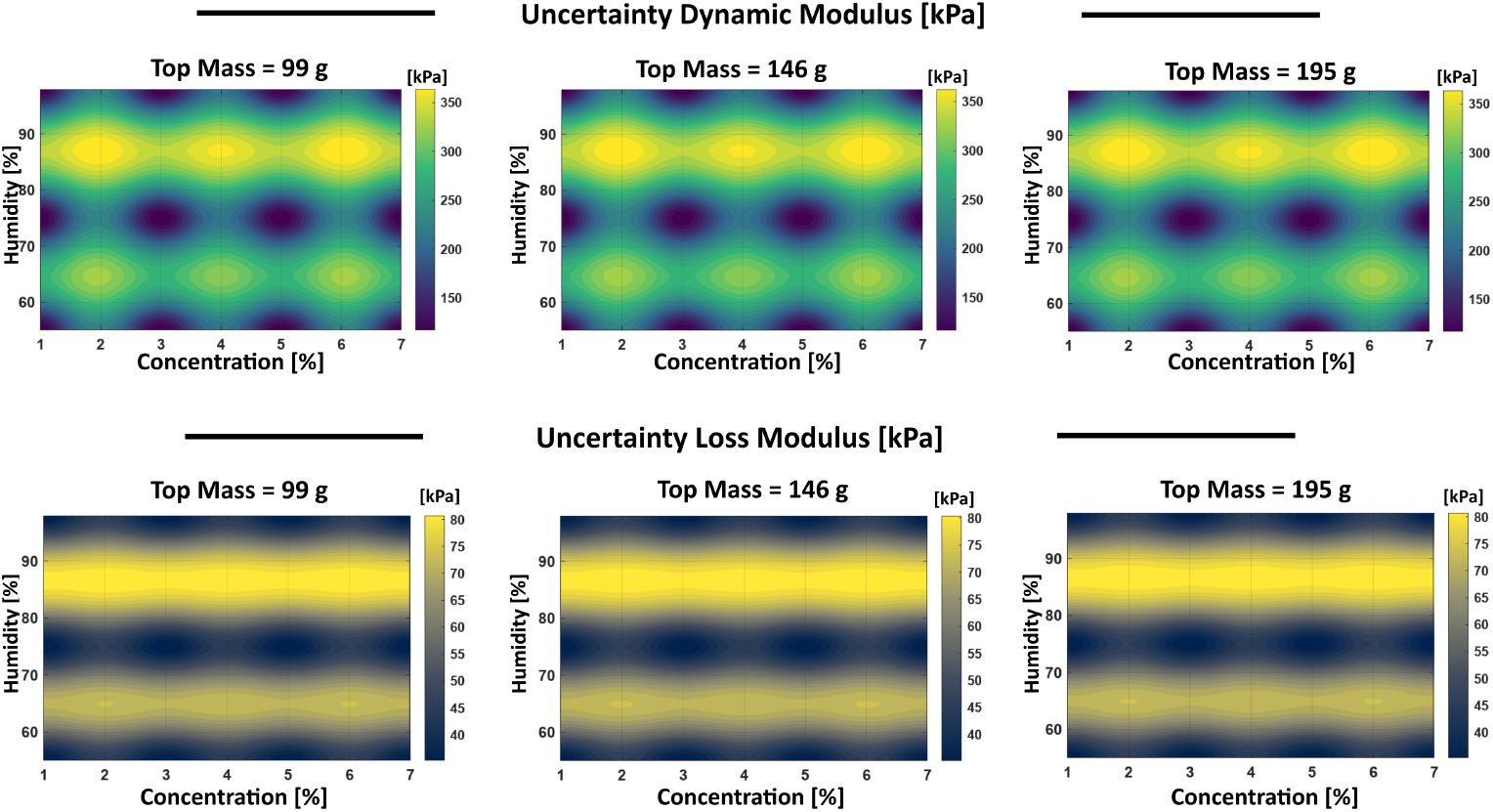
Predictive uncertainty maps of agarose gel viscoelastic properties. The heatmaps display the predictive standard deviation (*σ*) for the Dynamic Modulus (top row), and Loss Modulus (bottom row) as a function of Agarose Concentration and Relative Humidity. From left to right, the maps illustrate the model uncertainty at increasing pre-stress levels (Top Mass: 99 g, 146 g and 195 g). Darker regions (lower *σ*) correspond to the high-confidence zones surrounding experimental data points, while lighter regions (higher *σ*) represent interstitial predictive zones.

**Figure S3:**
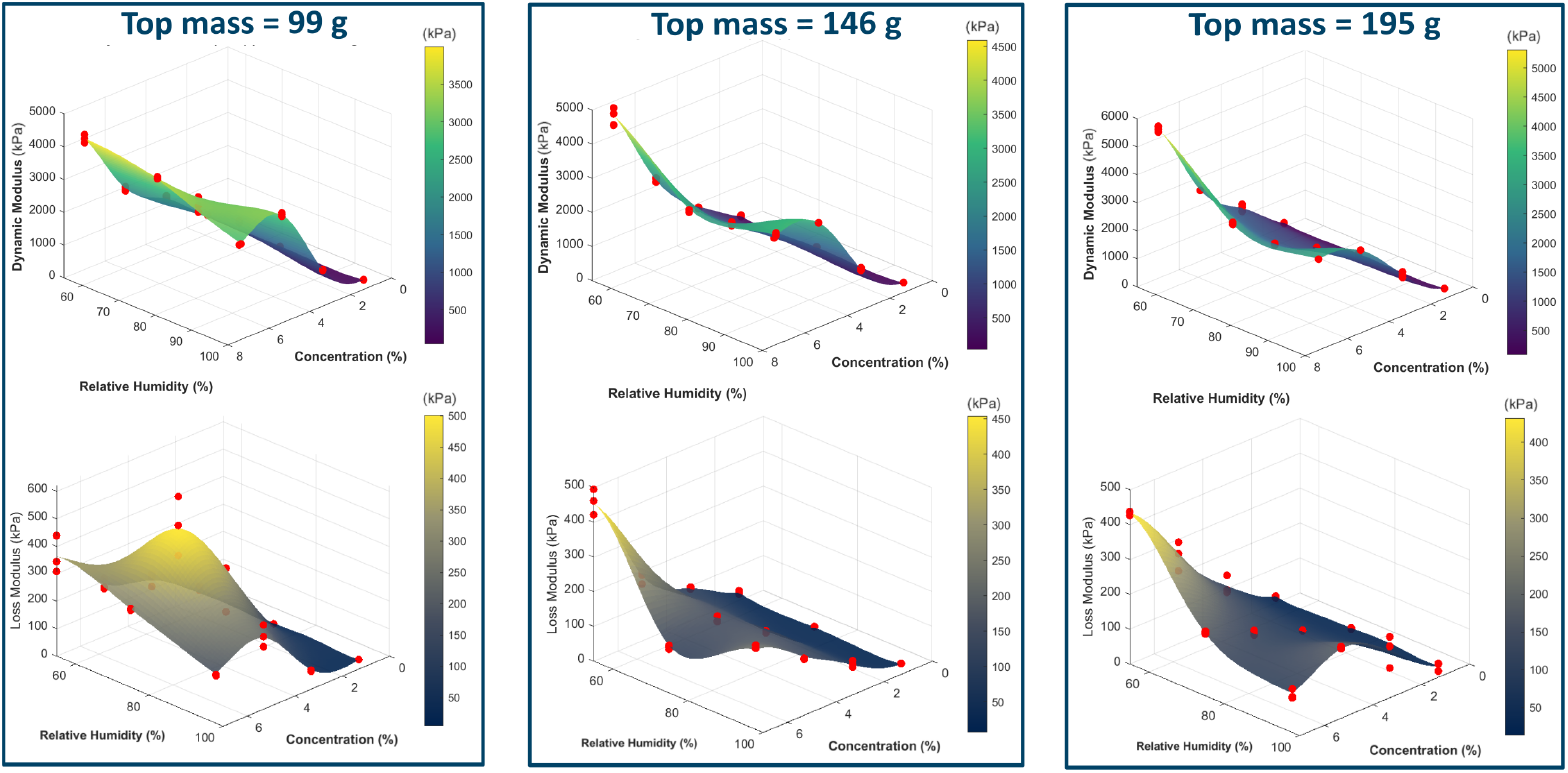
Gaussian Process Regression (GPR) response surfaces of agarose gel viscoelastic properties, experimental data points are overlaid as red spheres.

**Table S8:**
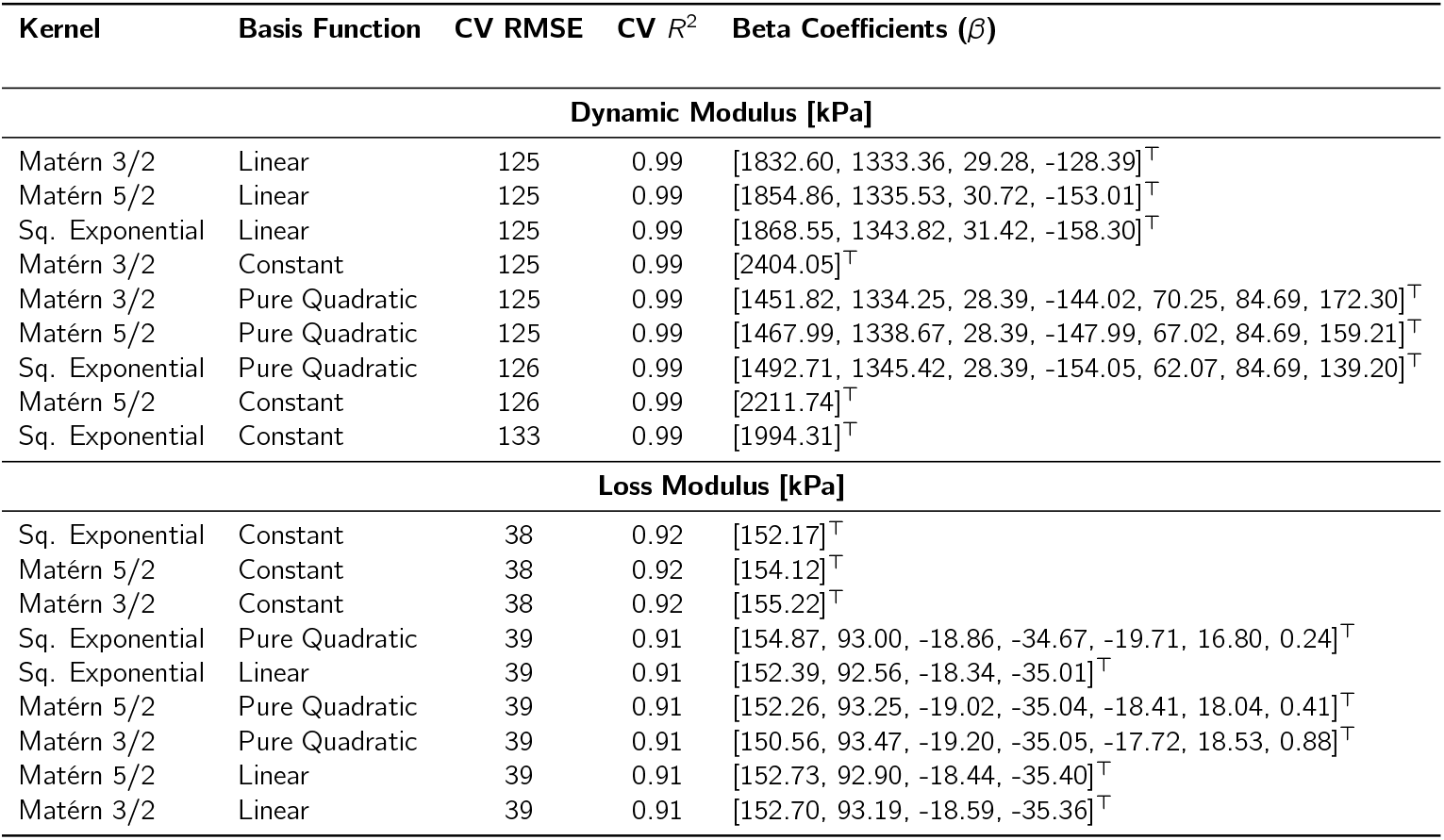
Evaluation of GPR architectures (kernel and basis function combinations) using grouped 3-fold cross-validation for predicting the viscoelastic properties of agarose gels. The table includes the optimized *β* coefficients.

**Figure S4:**
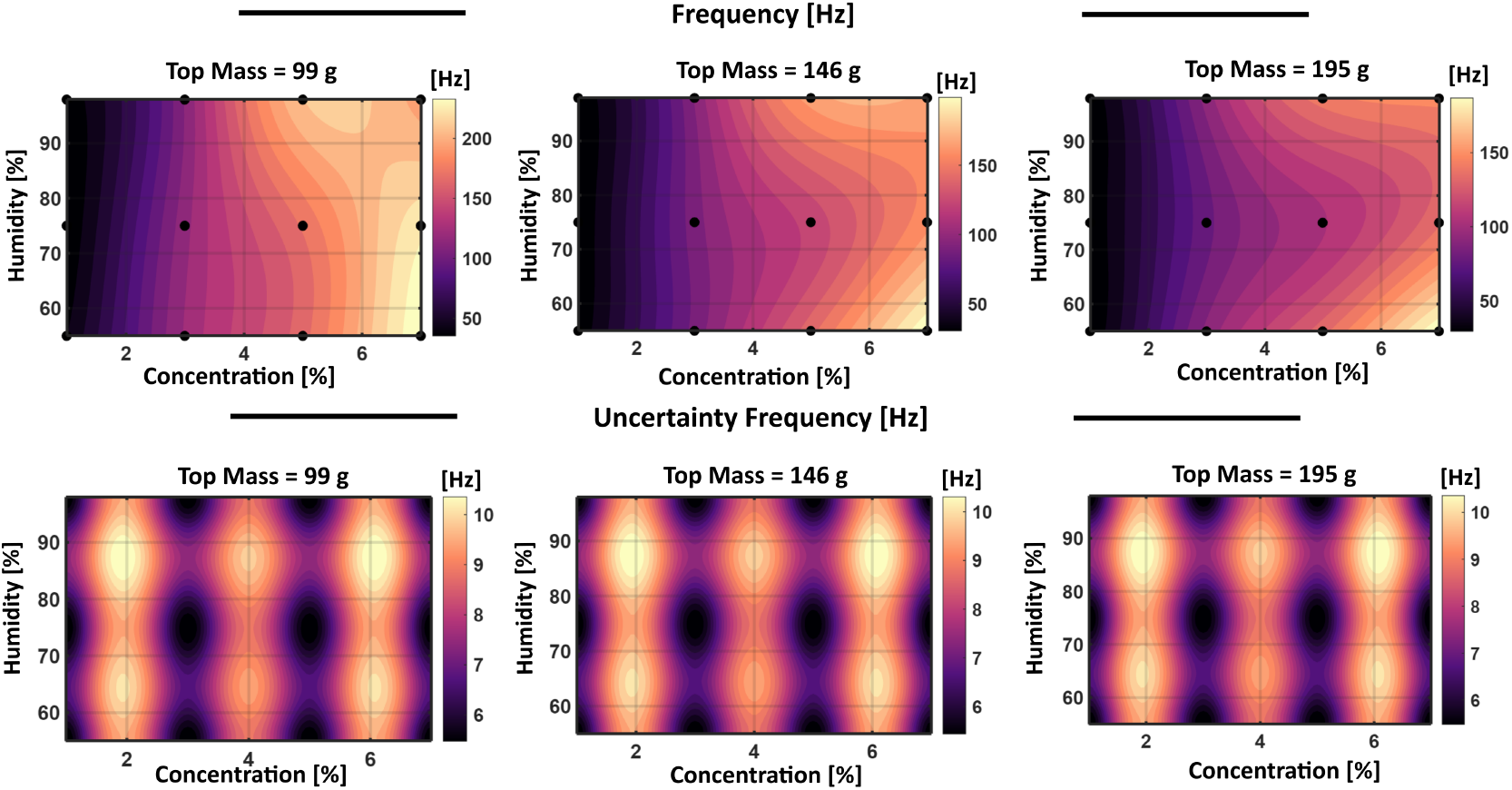
Gaussian Process Regression (GPR) contour maps of predicted resonance frequency (top) and associated model uncertainty (bottom) for agarose hydrogels. The response is evaluated as a function of polymer concentration and relative humidity under increasing compressive preloads (top mass). Black markers indicate the experimental sampling points. The model uses an Automatic Relevance Determination (ARD) `matérn` 5/2 kernel with constant basis set (CV RMSE = 5.88 Hz and 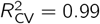)

**Table S9:**
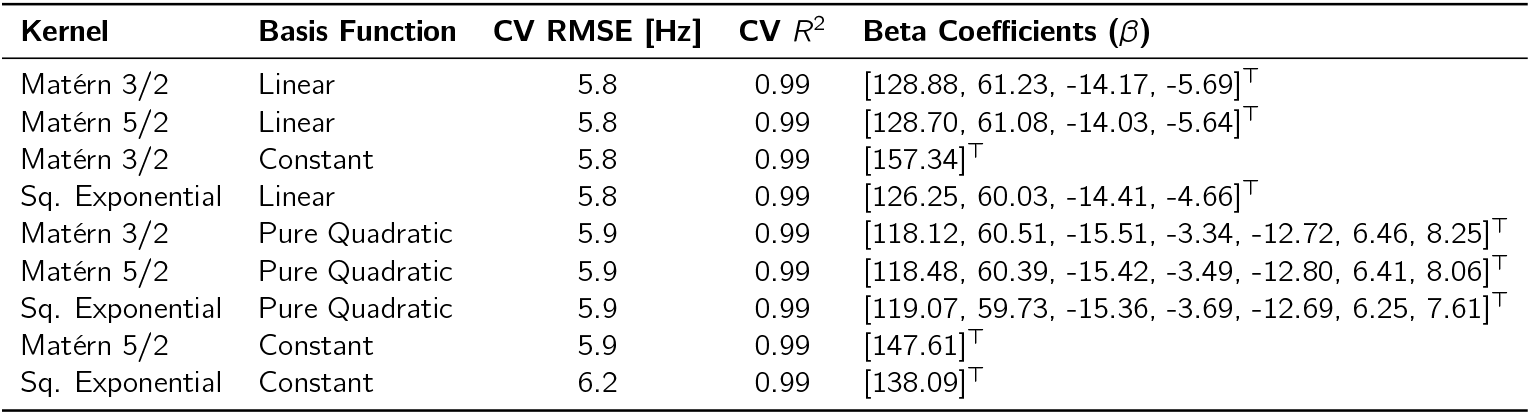
Evaluation of GPR architectures (kernel and basis function combinations) using grouped 3-fold cross-validation for predicting the resonance frequency of agarose hydrogels. The optimized *β* coefficients are reported for each model.

| Kernel | Basis Function | CV RMSE [Hz] | CV $R^2$ | Beta Coefficients ($\beta$) |
| --- | --- | --- | --- | --- |
| Matérn 3/2 | Linear | 5.8 | 0.99 | [128.88, 61.23, -14.17, -5.69]<sup>T</sup> |
| Matérn 5/2 | Linear | 5.8 | 0.99 | [128.70, 61.08, -14.03, -5.64]<sup>T</sup> |
| Matérn 3/2 | Constant | 5.8 | 0.99 | [157.34]<sup>T</sup> |
| Sq. Exponential | Linear | 5.8 | 0.99 | [126.25, 60.03, -14.41, -4.66]<sup>T</sup> |
| Matérn 3/2 | Pure Quadratic | 5.9 | 0.99 | [118.12, 60.51, -15.51, -3.34, -12.72, 6.46, 8.25]<sup>T</sup> |
| Matérn 5/2 | Pure Quadratic | 5.9 | 0.99 | [118.48, 60.39, -15.42, -3.49, -12.80, 6.41, 8.06]<sup>T</sup> |
| Sq. Exponential | Pure Quadratic | 5.9 | 0.99 | [119.07, 59.73, -15.36, -3.69, -12.69, 6.25, 7.61]<sup>T</sup> |
| Matérn 5/2 | Constant | 5.9 | 0.99 | [147.61]<sup>T</sup> |
| Sq. Exponential | Constant | 6.2 | 0.99 | [138.09]<sup>T</sup> |

**Figure S5:**
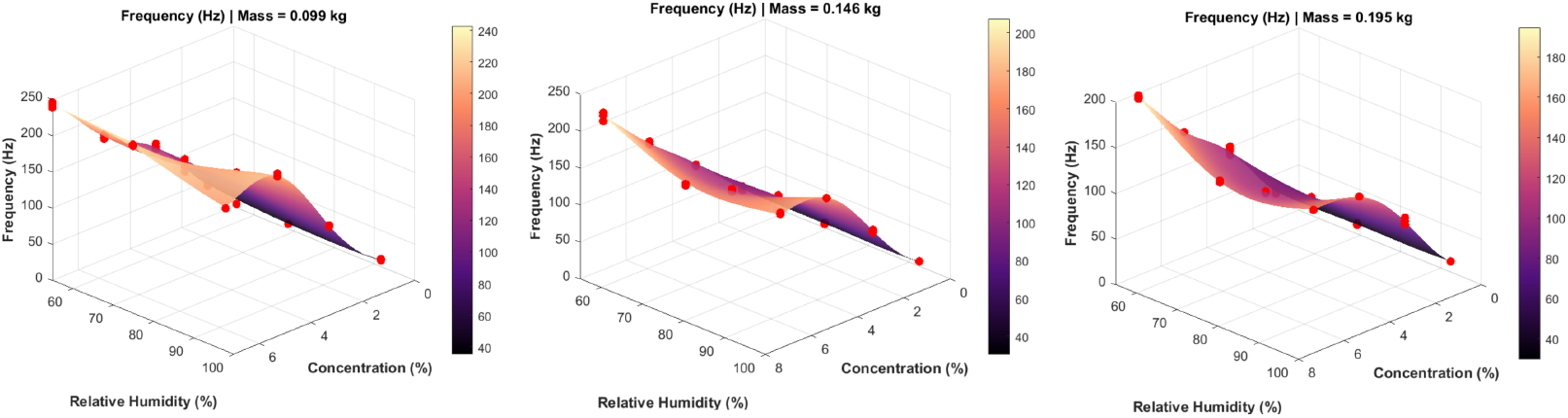
Gaussian Process Regression (GPR) response surfaces illustrating the effect of relative humidity and concentration on the frequency response of agarose gels under increasing preloads. Experimental data points are overlaid as red spheres.

## S6 VISCOELASTIC SENSITIVITY MAPS OF AGAROSE HYDROGELS

**Figure S6:**
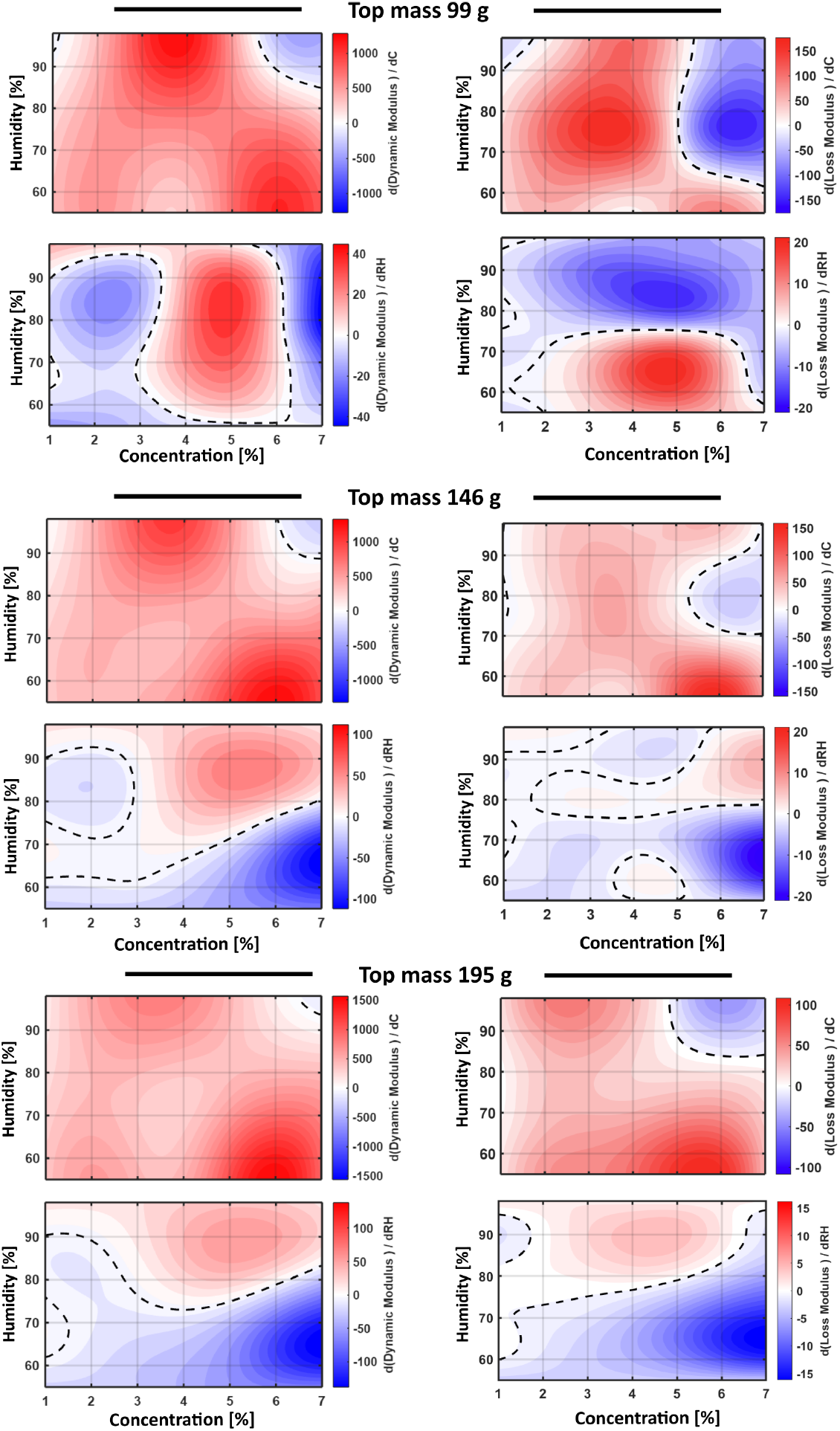
Viscoelastic sensitivity maps of agarose hydrogels evaluated under a different pre-stress mass from 99 g to 195 g. The contour plots visualize the partial derivatives of the material’s dynamic response with respect to polymer concentration. Dashed black lines represent zero-gradient contours, delineating critical structural and environmental transition zones where the material properties reach a local maximum, minimum, or plateau.

## S7 AGAROSE SAMPLE DATA: DYNAMIC MECHANICAL ANALYSIS

**Table S10:** Viscoelastic properties measured at a frequency of 28 Hz at 98% of humidity: Storage Modulus (*E*^*′*^) and loss modulus (*E*^*′′*^). Data are presented as Mean (Min – Max) for *n* = 3.

| Conc. | $E'$ [kPa] | $E''$ [kPa] |
| --- | --- | --- |
| 1% | 32 (26 - 40) | 5 (3 - 7) |
| 3% | 112 (110 - 114) | 13 (12 - 13) |
| 5% | 110 (61 - 192) | 13 (12 - 13) |
| 7% | 251 (170 - 378) | 20 (16 - 25) |

**Table S11:** Viscoelastic properties measured at a frequency of 28 Hz at 55% of humidity: Storage Modulus (*E*^*′*^) and loss modulus (*E*^*′′*^). Data are presented as Mean (Min - Max) for *n* = 3.

| Conc. | $E'$ [kPa] | $E''$ [kPa] |
| --- | --- | --- |
| 1% | 33 (15 - 71) | 8 ( 5 - 14) |
| 3% | 117 (82 - 176) | 10 (8 - 12) |
| 5% | 237 (113 - 368) | 20 ( 14 - 25) |
| 7% | 255 (133 - 337) | 22 (14 - 27) |

